# Divergent assembly trajectories: a comparison of the plant and soil microbiome with plant communities in a glacier forefield

**DOI:** 10.1101/2021.07.09.451772

**Authors:** Robert R. Junker, Xie He, Jan-Christoph Otto, Victoria Ruiz-Hernández, Maximilian Hanusch

## Abstract

Community assembly is a result of dispersal, abiotic and biotic characteristics of the habitat as well as stochasticity. A direct comparison between the assembly of microbial and ‘macrobial’ organisms is hampered by the sampling of these communities in different studies, at different sites, or on different scales. In a glacier forefield in the Austrian Alps, we recorded the soil and plant microbiome (bacteria and fungi) and plants that occurred in the same landscape and in close proximity in the same plots. We tested five predictions deduced from assembly processes and revealed deviating patterns of assembly in these community types. In short, microbes appeared to be less dispersal limited than plants, soil microbes and plants strongly responded to abiotic factors whereas the leaf microbiome was plant species-specific and well buffered from environmental conditions. The observed differences in community assembly processes may be attributed to the organisms’ dispersal abilities, the exposure of the habitats to airborne propagules, and habitat characteristics. The finding that assembly is conditional to the characteristics of the organisms, the habitat, and the spatial scale under consideration is thus central for our understanding about the establishment and the maintenance of biodiversity.

## Introduction

Species are heterogeneously distributed at global, regional, and local scales. Observed distributions are attributed to the species’ evolutionary history, dispersal abilities, adaptations to the environment, and interactions with other organisms as well as to drift as a stochastic element (Vellend, 2010). Species that share these characteristics and those that do not exclude each other, may co-occur more frequently than expected by chance and are thus often part of the same community (Götzenberger *et al.*, 2012). Depending on the scale and the organisms under consideration, dispersal filters, environmental filters, and / or interaction filters are the dominant processes explaining the composition and diversity of local plant or animal communities (Vellend, 2010, Vilmi *et al.*, 2020), which may be modulated by evolutionary and metacommunity dynamics (Mittelbach & Schemske, 2015). This line of research aiming at identifying the mechanisms underlying species co-occurrence and local diversity is central to ecological theory and nature conservation (HilleRisLambers *et al.*, 2012, Kraft *et al.*, 2015).

The increasing availability of data on bacterial and fungal communities has fueled the interest in microbial community assembly (Nemergut *et al.*, 2013). While the major processes in microbial community assembly are in principle the same as in ‘macrobial’ communities, striking differences between microorganisms and plant and animals may hamper a direct transfer of concepts and conclusions about the establishment and maintenance of diversity. Active dispersal in microbes is rare or restricted to very short distances for instance during active chemotaxis under ideal conditions (Raina *et al.*, 2019). On the other side, passive airborne dispersal may easily lead to intercontinental distributions of microbes because of smaller propagule sizes (Wilkinson *et al.*, 2012). High reproduction rates of microbes, high intraspecific genetic diversity, horizontal gene transfer, and rapid evolutionary responses to new habitats enable microorganisms to quickly occupy niches and consume resources (Nemergut *et al.*, 2013). Furthermore, and maybe most importantly, microbial community assembly occurs on different spatial scales compared to the assembly of plant and animals. Sampling a single leaf or petal means integrating over multiple microbial niches characterized by different availabilities of water, nutrients, and plant metabolites (Karamanoli *et al.*, 2012, Hayes *et al.*, 2021), whereas a one-square-meter-plot may be a rather homogenous niche for plants. Likewise, a soil particle is characterized by a strong gradient of abiotic variables and thus provides various niches (Sexstone *et al.*, 1985). Additionally, mutualistic and antagonistic interactions between microorganisms, which are a dominant factor in shaping microbial co-occurrence, occur on very small scales, often restricted to neighboring cells (Cordero & Datta, 2016, Dal Co *et al.*, 2020). Thus, field sampling protocols of plant and microbe communities address different organizational levels: a square meter represents a plant community of interacting species that occupying a largely uniform niche; a single leaf or soil particle hosts a number of microbial communities featuring separate interaction networks in diverse niches. Therefore, the relative importance of assembly processes may vary between plant, bacteria and fungi communities despite the fact that they colonize the same landscape. This, however, has not been directly compared.

The selection processes by which members of local communities are filtered from the regional species pool is uniformly considered to be mostly determined by dispersal, abiotic conditions (environment) and species interactions (Fig. 1) (de Bello *et al.*, 2012, Götzenberger *et al.*, 2012, Cadotte & Tucker, 2017), which is basically also covered in Vellend’s (2010) conceptual synthesis. The dispersal filter assumes variation in the dispersal abilities of the species present in the regional species pool, which leads to different sets of species that reach a given location. One prediction of the dispersal filter hypothesis is that communities are more similar to each other when they are located in close proximity, meaning that their dissimilarity increases with larger distances between the communities (Fig. 1, H1). Such a pattern was detected in some microbial systems but not in others (Belisle *et al.*, 2012, Donald *et al.*, 2020) but the factors explaining these contrasting results remain unknown. Once organisms reached a given habitat, the environmental filter hypothesis states that abiotic conditions determine whether a species is able to survive and reproduce. This hypothesis predicts that habitats with similar abiotic conditions host more similar communities than habitats that differ in abiotic variables (Fig. 1, H2). The environmental filter hypothesis is well supported for a number of microbial systems (Berg & Smalla, 2009, Bulgarelli *et al.*, 2013). More specifically for plant-associated microbes, the plant phenotype can be regarded as environment for microbial communities leading to the prediction that plant species host specific microbial communities (Fig. 1, H3). This hypothesis has also been verified in numerous studies (Laforest-Lapointe *et al.*, 2016, Gaube *et al.*, 2021) suggesting that plant species-specific properties control microbial colonization (Junker & Tholl, 2013, Junker & Keller, 2015, Boachon *et al.*, 2019). Finally, even in suitable environments, resident species may prevent a successful establishment of further species due to competitive or inhibitory effects. Alternatively, species may also facilitate their establishment. Both of these processes are summarized as interaction filter. In the case of antagonistic interactions, the interaction filter hypothesis would predict lower co-occurrences of species than expected by chance; in case of mutualistic interactions higher co-occurrences than expected by chance (Fig. 1, H4). Interactions between microbes can be strong (Cordero & Datta, 2016, Dal Co *et al.*, 2020), but whether these interactions contribute to community establishment in microbes remains understudied (Nemergut *et al.*, 2013). In this study, we test a fifth hypothesis specific to above-ground plant-associated microbial communities, which does not directly address the assembly process of communities, but rather the source for these microbes. It has been suggested that soil is a reservoir for bacteria associated with roots and also above-ground plant organs (Berg & Smalla, 2009, Bulgarelli *et al.*, 2013, Bai *et al.*, 2015). Thus, these findings suggest that leaf-associated microbial communities consist of a subset of the microbes found in the soil plus those specific to above-ground plant parts (Fig. 1, H5). Using data on community composition to infer assembly processes has been criticized (Gilbert & Bennett, 2010, Stegen & Hurlbert, 2011, Kraft *et al.*, 2015, Stegen *et al.*, 2015, Cadotte & Tucker, 2017, Blanchet *et al.*, 2020) and our tests are not meant to identify the dominant assembly process for the communities under consideration. However, our comparative approach considering communities composed of different organisms that colonize different habitats is well suited to reveal fundamental differences between the ecology and assembly of these communities.

**Fig. 1.**
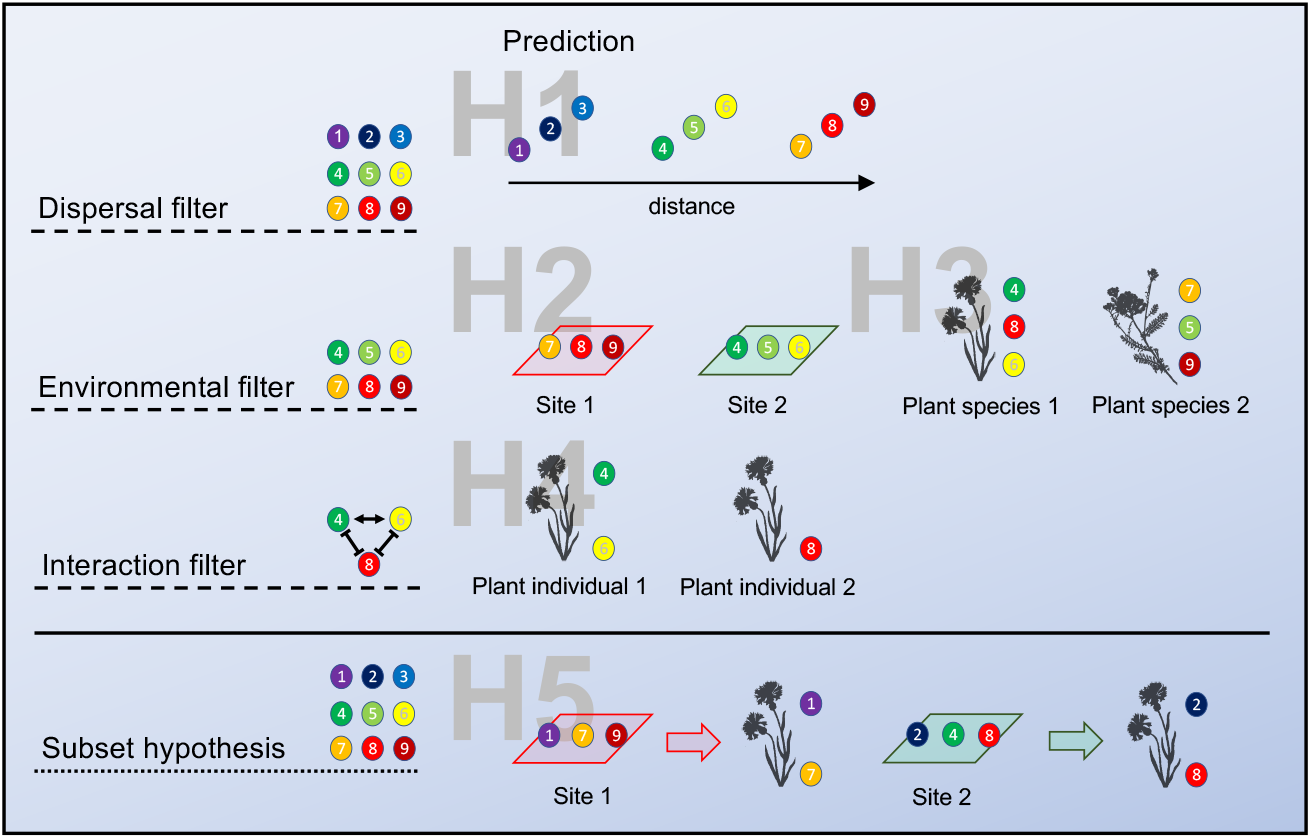
Hypotheses on the assembly of communities. Each circle (1-9) represents an OTU or species. Hypotheses H1, H2, and H4 are adopted from classical macroscopic community ecology: The dispersal filter (H1) selects species from the regional species pool that are able to reach the habitat. Once the species entered a given habitat, the environmental filter (H2) selects those species that are able to establish and reproduce given the local abiotic conditions. Finally, the interactions filter (H4) select species that are either facilitated or are at least not outcompeted by the resident species, i.e. a community consists of taxa that have the potential to co-occur. Hypotheses H3 and H5 are more specific to above-ground plant-associated bacteria or fungi. Classically, the environmental filter addresses abiotic parameters such as temperature or pH. The plant itself is, however, the habitat for bacteria and fungi that colonize above-ground plant parts (H3). The properties of these habitats are plant-organ but also plant-species specific, thus plant-species identity is a good proxy for the conditions on the plant habitat. The subset hypotheses (H5) is more about the origin of the microbes than on the assembly. It states that the soil is an important pool for microbes associated with above-ground plant organs. Their establishment on above-ground plant organs is again dependent on assembly processes as described above.

Community assembly processes of bacteria and fungi have been studied either focusing on microbes inhabiting soil or associated with leaves (Schmidt *et al.*, 2014, Donald *et al.*, 2020, Gao *et al.*, 2020, Hassani *et al.*, 2020). We estimated the importance of five assembly hypotheses (Fig. 1) for bacterial and fungal communities found in soil and on leaves as well as for plant communities in a comparative approach using null models. These five community types were sampled in the same *n* = 140 plots along a successional gradient in a glacier forefield in the Austrian Alps, which allows a direct comparison of how the different assembly processes affect different communities within the same landscape. Our study reveals detailed insights into the dispersal abilities of microbes and the biotic and abiotic factors shaping bacterial and fungal communities in comparison to plants and thus contributes to our understanding about the establishment and the maintenance of biodiversity.

## Material and Methods

### Study site

The study was conducted in the long-term ecological research platform Ödenwinkel which was established in 2019 in the Hohe Tauern National Park, Austria (Dynamic Ecological Information Management System – site and dataset registry: https://deims.org/activity/fefd07db-2f16-46eb-8883-f10fbc9d13a3, last access: March 2021). A total of *n* = 140 permanent plots was established in the valley of the Ödenwinkelkees glacier. 135 plots are located within the glacier forefield (from the glacier mouth: plots 1 to 135), which were covered by ice at the latest glacial maximum in the Little Ice Age (LIA; around 1850). The remaining plots (plots 136 to 140), were established in areas outside the glacier forefield. Plots within the glacier forefield were evenly distributed, representing a successional gradient spanning over 1.7km in length. Plots were defined as squares with an area of 1 m^2^ and were all oriented in the same cardinal direction. Further details on the design of the research platform, exact plot positions, as well as on the sampling strategy can be found in Junker et al. (2020). During field season in 2019, we estimated the abundance of all vascular plant species growing on the plots and installed a temperature logger (MF1921G iButton, Fuchs Elektronik, Weinheim, Germany) 10 cm north of each plot centre, at a depth of 3 cm below ground and calculated the mean seasonal temperature which has been shown to affect plant species composition as well as interactions between plants and other organisms (Ohler *et al.*, 2020). The thermo loggers were set to start on 13^th^ August 2019 and were stopped on 9^th^ August 2020 with a total of 2048 measurements recorded over 362 days. Mean seasonal temperature was calculated on the basis of the recordings ranging from 26^th^ of June to 16^th^ of September representing the period in which the plots were free of permanent snow cover before and after the winter 2019/2020. Exact coordinates of each plot were directly exported from a GPS device and the distance to closest stream was retrieved from a digital elevation model (1 mLIDAR DEM, Land Salzburg, see Junker *et al.*, 2020). In 2020, soil samples were taken and soil nutrients (Ca, P, K, Mg and total N2) as well as soil pH were measured on all plots (except for plot 129) by AGROLAB Agrar und Umwelt GmbH (Landshut, Germany).

### Sampling of microbiome

We sampled microorganisms (bacteria and fungi) inhabiting the phyllosphere of the most frequently occurring plant species and the soil of each plot. Sampling was performed within 11 days during the main vegetative period (July 31^th^ – August 10^th^ 2019). Leaf and soil samples were collected using sterilized forceps (dipped into 70 % ethanol and flamed) to avoid contamination. We sampled bacterial and fungal communities in the phyllospheres of three focus plant species on every plot where they occurred: *Oxyria digyna* as representative of early succession, *Trifolium badium* as representative of late succession, and *Campanula scheuchzeri* which occurred all along the successional gradient (for detailed information on the selection of the focus plant species see Junker *et al.*, 2020). Furthermore, we took three samples of the most frequently found vascular plant species, i.e. species that occurred on 10 or more plots (*n* = 45 species). In these cases, we took samples on the oldest, the youngest, and the intermediate plot where they occurred. For every plant sample, we took 1 to 3 leaves according to different leaf sizes of the species to make sure that the size of the leaf samples was largely consistent among species. Soil microbiome samples were taken as pooled samples from two locations on every plot whenever there was enough soil to proceed. With a bulb-planting device, we took soil cores, from which we took soil samples at 3 cm depth. In plots where it was not possible to take soil cores due to a lack of developed soil, we collected sediment underneath or next to rocks. Collected samples were directly transferred to ZR BashingBeads Lysis tubes containing 750 μL of ZymoBIOMICS lysis solution (Zymo-BIOMICS DNA Miniprep Kit; Zymo Research, Irvine, California, USA). Within 8 h after collection of microbial samples, ZR BashingBeads Lysis tubes were sonicated for 7 min to detach microorganisms from the surfaces. In the case of plant leaves, we removed them from tubes next to a flame with sterile forceps after the sonication to decrease the amount of plant DNA in the samples. Subsequently, all microbial samples were shaken using a ball mill. In cases where we were able to fully remove plant tissues from collection tubes and soil samples, tubes were shaken for 9 min with a frequency of 30.0 s^-1^. In some cases, it was not possible to fully remove plant tissues from tubes, and samples were shaken for 5 min at 20.0 s^-1^. Microbial DNA was extracted using the ZymoBIOMICS DNA Miniprep Kit following the manufacturer’s instructions. Next-generation sequencing and microbiome profiling of isolated DNA samples were performed by Eurofins Genomics (Ebersberg, Germany, for more information see Junker *et al.*, 2020).

### Test of hypotheses on community assembly

To test the hypotheses on community assembly for specific groups of microbes, we generated the following subsets of the datasets on bacterial and fungal communities: bacteria and fungi a) associated with plants (bacteria: *n* = 308; fungi: *n* = 324), b) colonizing the soil (bacteria: *n* = 132; fungi: *n* = 135), and c) associated with three focus plant species (*Campanula scheuchzeri* (bacteria: *n* = 94; fungi: *n* = 113), *Oxyria digyna* (bacteria: *n* = 19; fungi: *n* = 26), *Trifolium badium* (bacteria: *n* = 50; fungi: *n* = 23)). In total, we recorded the composition of *n* = 140 plant communities. In the following, the different subsets are referred to as ‘community types’. Prior to the statistical analysis of microbial communities, we performed a cumulative sum scaling (CSS) normalization (R package *metagenomeSeq* v1.28.2) on the count data to account for differences in sequencing depth among samples. Additionally, to compare the assembly processes of bacteria and fungi to those of plants, we also used plant cover as a surrogate of abundance recorded at all of the *n* = 140 plots. To test the assembly hypotheses, we first performed the statistical analyses as described below using the field data, and additionally for null models generated from the same data. For each subset of the data we generated *n* = 1000 null models using the function *nullmodel, method = “r2d”*implemented in the R package *bipartite* (Dormann *et al.*, 2014). This method generates random community tables with fixed row and column sums using the Patefield’s algorithm (Patefield, 1981). For each hypothesis and data subset, we generated one test statistic for the field dataset and *n* = 1000 test statistics for the null models generated from the field data. As a measure of deviation of the observed result from the null model expectation, we used one-sample Cohen’s *d* = (*Mean - Mu*) / *Sd* with *Mean* and *SD* as the mean value and standard deviation of null model results and *Mu* as observed result. Higher Cohen’s *d* values indicate stronger effect. As Cohen’s *d* thus is based on different test statistics with different ranges (see below) the values are not comparable between the hypotheses, but provide a good measure to compare the effects on different groups (bacteria, fungi detected on leaves or soil, plants) within a hypothesis. Significant differences between null models and observed results were indicated if the observed result did not overlap with the 95% confidence interval of results obtained from null models.

### H1: dispersal filter

If dispersal filter determined the composition of communities, we would expect that communities spatially close to each other are more similar to each other in their composition than communities that are separated by larger distances. As spatial distance we used Euclidean distances based on the latitude, longitude and elevation of the plots where we sampled the communities. For the similarity in community composition, we used Bray-Curtis distances based on the CSS abundance of the OTUs in the case of bacteria and fungi or the abundance of plants. To test for a correlation between the spatial distance and community distance we performed Mantel test based on Pearson’s product-moment correlation, the *r*-value was used as test statistic. In the analysis using all samples of leaf associated microbes, potential effects of other assembly rules (niche-based processes) may overlay the effect of dispersal limitation. Therefore, we also performed the analysis only within the samples collected from leaves of one of the three focus species, which represent a more uniform habitat.

### H2 and H3: environmental filter

If environmental filters determined the composition of communities, we would expect that communities established on plots characterized by similar environmental parameters would be more similar to each other compared to communities established in different environments. As environmental distance (H2) we used Euclidean distances based on the soil nutrients (N, P, K, Mg), soil pH, distance to closest stream, and the mean seasonal temperature. Additionally, we used the plant species composition as environmental parameter for microbes and used Bray-Curtis distances to calculate the distances between plots. For the similarity in community composition, we used Bray-Curtis distances based on the CSS abundance of the OTUs in the case of bacteria and fungi or the abundance of plants. To test for a correlation between the environmental distance and community distance we performed Mantel statistic based on Pearson’s product-moment correlation, the *r*-value was used as test statistic. Again, for leaf-associated bacteria and fungi, we repeated this analysis using only the samples collected from leaves of one of the three focus species. For leaf associated bacteria and fungi, not necessarily the soil parameters define the environmental niche, but the physical and chemical properties of the leaves. Therefore, to test hypothesis 3, we performed distance-based redundancy analyses using Bray-Curtis distances followed by ‘permutation test under reduced model’ to test whether bacterial and fungal communities are more similar within than between plant species. The *F*-values of permutation test was used as test statistic. We did this for the whole dataset comprising all plant species sampled and also for a subset considering only the three focus species that have a meaningful sample size.

### H4: interaction filter

If interaction filters determined the composition of communities, we would expect that OTUs show a higher co-occurrence (higher aggregation) than expected by chance if facilitation between species is the dominant type of interaction; or show a lower co-occurrence (higher segregation) than expected by chance if competition between species is the dominant type of interaction. To test for species aggregation or segregation we used the *cooc_null_model* function implemented in the R package *EcoSimR.* The C-score was used as metric to evaluate whether cooccurrence patterns are rather aggregated (low C-scores) or segregated (high C-scores). For this hypothesis, we only used data sets on bacteria and fungi associated to leaves of the three focus species, bacteria and fungi in soil, and plant communities to make sure that the organisms share a common habitat where they can interact.

### H5: subset hypothesis

If soil was a major source of plant associated bacteria and fungi, we would expect that the proportion of leaf-associated OTUs that are found in both leaf and soil samples is higher when tested using soil samples from the same plot were the leaf was sampled as compared to soil samples from other plots. Thus, for each plant sample we first calculated the proportional overlap between leaf-associated OTUs and the OTUs detected in the soil of the plot were the plant was sampled. As a second step, we calculated the proportional overlap between leaf-associated OTUs and the OTUs detected in the soil sampled in all other plots. Therefore, here the second step represent the null model. The mean overlap of leaf-associated microbes with soil microbes on the same plot was used as *Mu*; the mean and standard deviation of the mean overlaps of leaf-associated microbes with soil microbes on different plots as *Mean* and *SD* in the formula to calculate Cohen’s *d*.

## Results

In total, we detected *n* = 10860 bacterial OTUs associated with plant leaves, *n* = 5221 bacterial OTUs in soil samples, *n* = 5363 fungal OTUs associated with plant leaves, *n* = 6014 fungal OTUs in soil samples, and *n* = 108 plant species. Raw sequences of next-generation 16S rRNA gene amplicon sequencing are available at the NCBI Sequence Read Archive (SRA) under the BioProject accession PRJNA701884 and PRJNA701890.

### H1: dispersal filter

We did find only a weak correlation between the similarity of bacterial and fungal communities associated with leaves and the spatial distance between these communities (Fig. 2a, Supplementary Information 1). In contrast, microbial soil communities as well as plant communities showed a strong correlation between their composition and spatial distance, i.e. communities in close proximity showed a higher similarity in composition than communities separated by larger distances (Fig. 2a, Supplementary Information 1). In all cases the observed Mantel *r* was larger than the Mantel *r* expected from null models. Microbial communities associated with *Oxyria digyna* and *Trifolium badium* also did not show a strong correlation between compositional similarity and spatial distance (Fig. 2b, Supplementary Information 1). However, communities associated with *C. scheuchzeri* leaves, the plant species with the largest range of distribution within the successional gradient, showed a moderate relationship between compositional similarity and spatial distance (Fig. 2b, Supplementary Information 1) whereas null model expectations were close to zero for Mantel *r*.

**Fig. 2.**
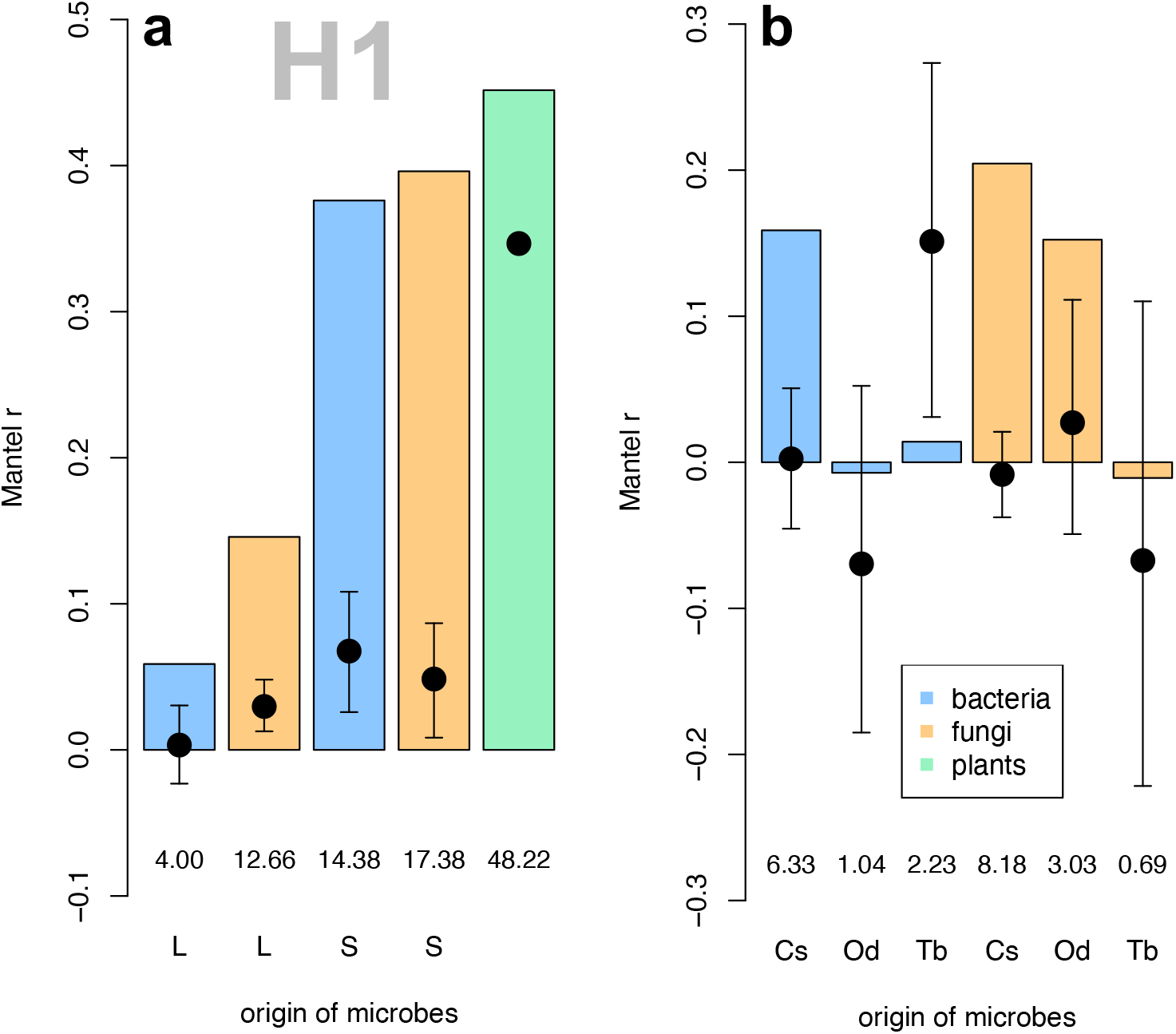
Correlations between the community similarity and the spatial distance between the communities considering bacteria (blue bars) and fungi (orange bars) associated with leaves of all species (**a**, marked with L below the bars) and with leaves of one of the three focus plant species (**b**, marked with Cs for *Campanula scheuchzeri,* Od for *Oxyria digyna,* or Tb for *Trifolium badium* below the bars), or soil bacteria and fungi (**a**, marked with S below the bars), or plant species (**a**, green bar). Bars denote observed Mantel *r*-values, the circles denote Mantel *r*-values from null model expectations with 95% confidence intervals. The numbers below bars denote effect size Cohen’s *d.*

### H2: environmental filter

To test the environmental filter hypothesis, we used two datasets to characterize the environment of microbes and plants: 1) abiotic factors: soil nutrients (N, P, K, Mg), soil pH, distance to closes stream, and the mean seasonal temperature (Fig. 3a, b); 2) the biotic environment, which is the composition of plant species growing in each plot (Fig. 3c, d). Similarity in microbial community compositions associated with leaves showed no or weak correlations with the similarity in abiotic and biotic characteristics of the plots, both considering all samples of bacteria and fungi associated to plants (Fig. 3a, c, Supplementary Information 1) or those associated with one of the focus species (Fig. 3b, d, Supplementary Information 1). As an exception, communities of leaf associated fungi showed moderate responses to the biotic environment of the plots (Fig. 3d). In contrast, communities of soil microbes and plants found in plots with similar abiotic and biotic characteristics where clearly more similar in their composition compared to those communities found in different environments (Fig. 3a, c, Supplementary Information 1). Abiotic environmental factors were more similar between plots in close proximity compared to plots in larger distances (Mantel statistic based on Pearson’s product-moment correlation: *r* = 0.32, *p* = 0.001).

**Fig. 3.**
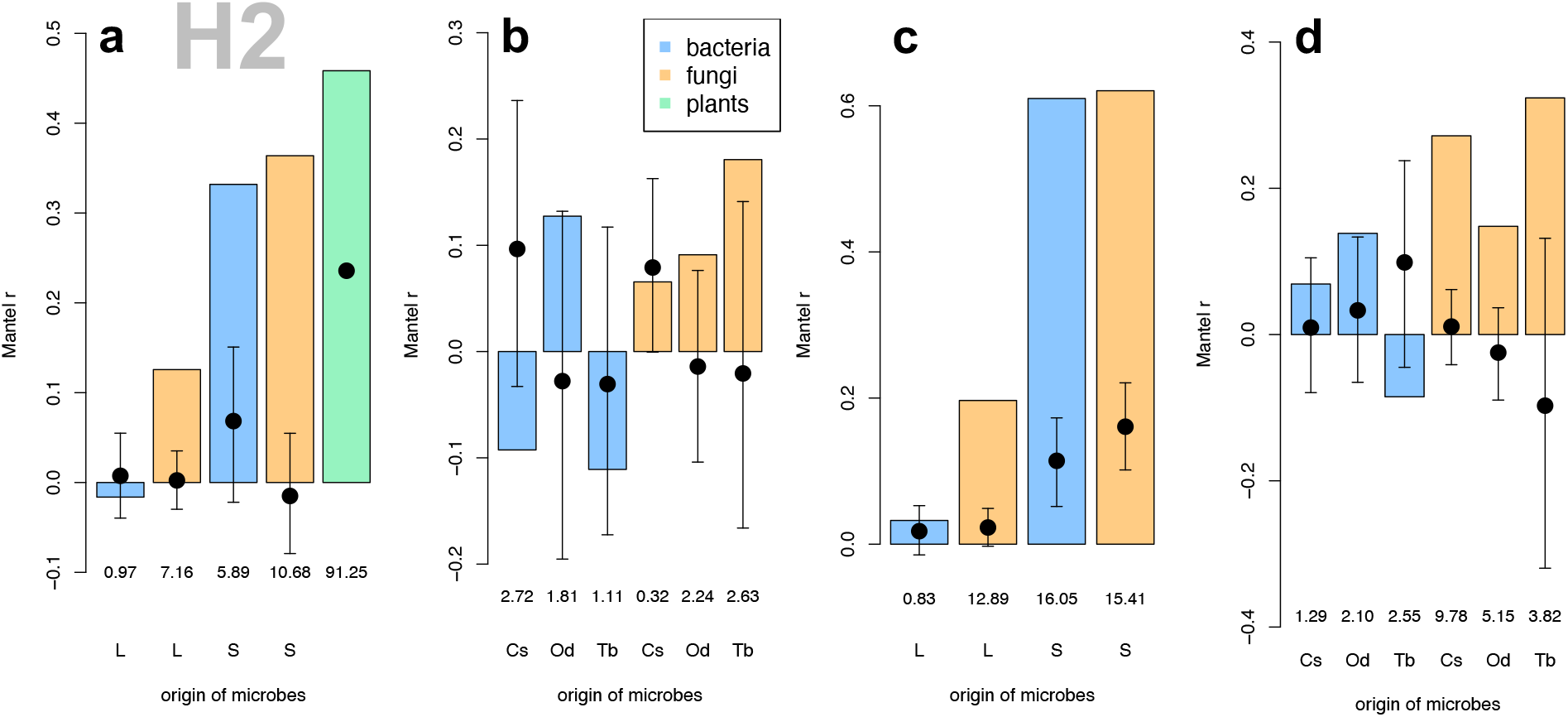
Correlations between the community similarity and the similarity of environmental parameters based on abiotic properties (**a**, **b**) or plant species composition (**c**, **d**) considering bacteria (blue bars) and fungi (orange bars) associated with leaves of all species (**a** and **c**, marked with L below the bars) and with leaves of one of the three focus plant species (**b** and **d**, marked with Cs for *Campanula scheuchzeri,* Od for *Oxyria digyna,* or Tb for *Trifolium badium* below the bars), or soil bacteria and fungi (**a** and **c**, marked with S below the bars), or plant species (**a**, green bar). Bars denote observed Mantel *r*-values, the circles denote Mantel *r*-values from null model expectations with 95% confidence intervals. The numbers below bars denote effect size Cohen’s *d.*

### H3: environmental filter

Plant species identity turned out to be a strong predictor for bacterial and fungal communities associated with leaves (Fig. 4, Supplementary Information 1), an effect that was even more pronounced when the distance-based redundancy analyses were restricted to the three focus plant species (Fig. 4b, c).

**Fig. 4.**
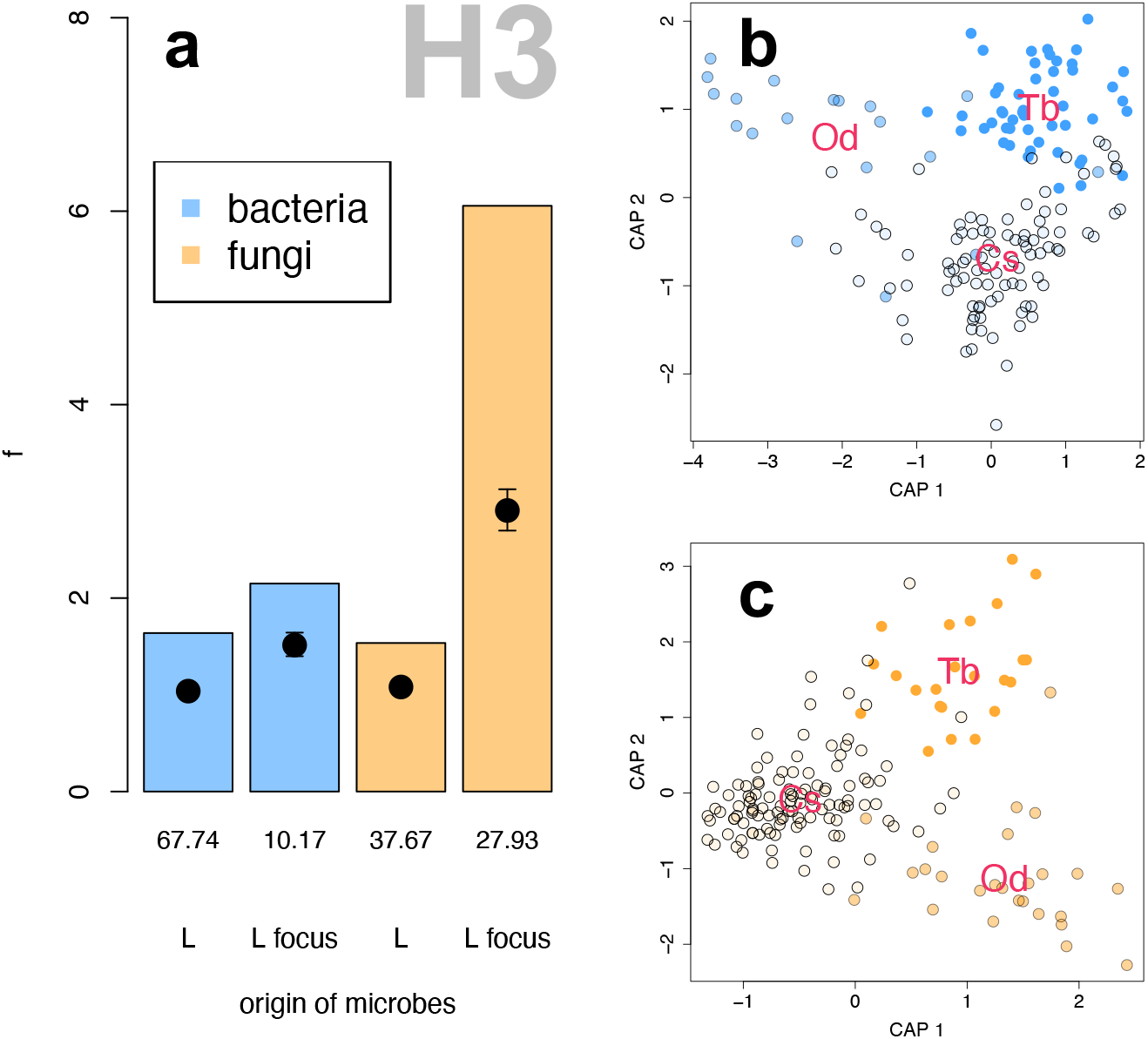
Composition of microbial OTUs associated with leaves is explained by plant species identity. Bars denote observed *f*-values of distance-based redundancy analyses using Bray-Curtis distances followed by permutation test under reduced model, the circles denote *f*-values from null model expectations with 95% confidence intervals. The numbers below bars denote effect size Cohen’s *d* (**a**). Either all leaf samples of bacteria (blue bars) and fungi (orange bars) are consideres (marked with L below the bars) or only samples of the three focus plant species (marked with L focus below the bars). Ordination based on distance-based redundancy analyses using Bray-Curtis distances of bacterial (**b**) and fungal (**c**) communities associated with leaves of the three focus plant species. Centroids of the three communities are indicated by Cs for *Campanula scheuchzeri* (very pale colors, black frame), Od for *Oxyria digyna* (pale colors, grey frame), or Tb for *Trifolium badium* (saturated colors, no frame).

### H4: interaction filter

Observed mean co-occurrence of OTU / species pairs was lower than null model expectations in all communities tested, i.e. observed C-scores were higher than C-scores obtained from null models (Fi. 5a, Supplementary Information 1). Overall, segregation of bacterial and fungal OTU pairs was higher in soil samples than in most leaf samples. Plants showed strongest segregation.

### H5: subset hypothesis

The proportion of leaf-associated OTUs that are found in both leaf and soil samples was higher when leaf and soil samples originated from the same plot than the mean proportion of shared OTUs when leaf and soil samples did not originate from the same plot (null model expectation, Fig. 5, Supplementary Information 1). However, observed proportion of overlapping bacterial and fungal OTUs was within the 95% confidence interval from null model expectation.

**Fig. 5.**
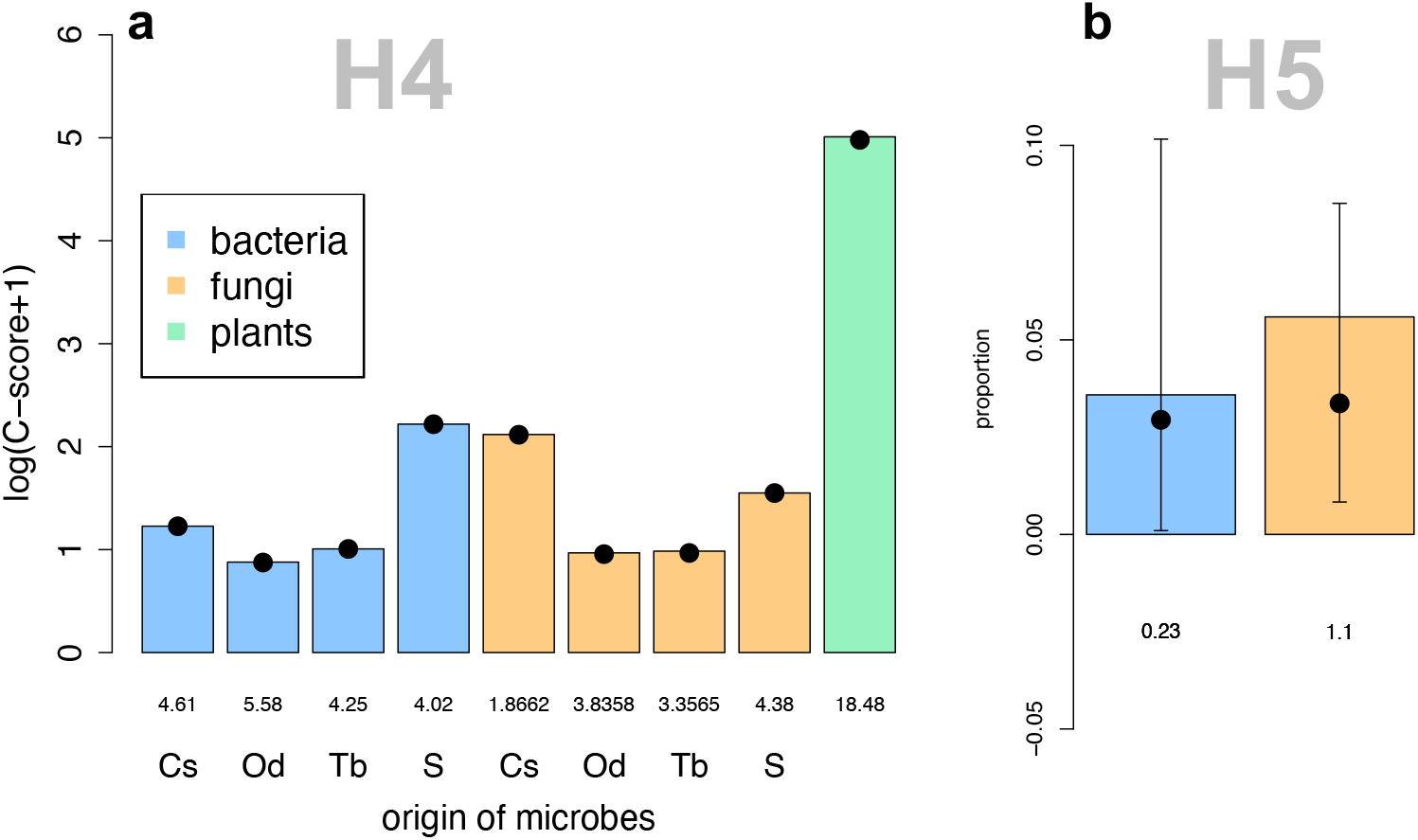
Mean co-occurrence between pairs of bacterial OTUs (blue bars) and pairs of fungal OTUs (orange bars) associated with leaves of the three focus plant species (**a**, marked with Cs for *Campanula scheuchzeri*, Od for *Oxyria digyna*, or Tb for *Trifolium badium* below the bars), or found in soil (**a**, marked with S below the bars), or plant species (green bar). Bars denote observed *C-scores,* the circles denote *C-scores* from null model expectations with 95% confidence intervals (note that 95% confidence intervals are too small to be visible). C-scores of null models were lower than observed C-scores in all cases. The numbers below bars denote effect size (**a**). Mean proportional overlap between leaf and soil OTUs (**b**). Bars denote mean observed proportion of OTUs that are found on the leaf and the soil of the same plot, the circles denote the proportional overlap from null model expectations with 95% confidence intervals. The numbers below bars denote effect size Cohen’s *d* (**b**).

## Discussion

The assembly of local communities is shaped by the interplay of different mechanisms that determine the occurrence, co-occurrence and diversity of species. In our approach, we operationalized the individual processes that contribute to community assembly by deducing specific predictions and testing them on datasets on five community types sampled in the same landscape: bacterial and fungal communities colonizing soil or are associated with leaves, and plant communities. Our results show that plant communities contain the strongest spatial signal, followed by microbes colonizing the soil; similarity of plant-associated bacterial and fungal communities was independent of spatial distance. Likewise, plant and soil microbe community compositions strongly responded to the environment whereas plant-associated microbes did not or only weakly. However, we found plant species-specific microbial communities associated with the leaves, supporting the notion that leaf characteristics constitute the environmental conditions for these microbes (Junker & Tholl, 2013). The observed co-occurrence patterns deviated slightly positively from null model expectations in all community types indicating that these patterns are either the result of random species distributions or that antagonistic interactions led to the segregation of species. Finally, we identified some bacterial and fungal OTUs that occurred in both soil and leaf samples, but the proportion of leaf-associated OTUs that were also detected in soil samples was low and was not higher in cases when the leaf and soil sample originated from the same plot. These results suggest that soil microbial communities and plant communities are shaped by dispersal limitation and / or environmental filtering. In contrast, leaf-associated microbial communities are not dispersal limited and are largely buffered from environmental conditions; instead leaf characteristics replace environmental parameters and strongly affect the community composition of bacteria and fungi associated with leaves. The interaction filter seemed to be relaxed for all community types in our study area.

In principle, microbes are less dispersal limited than ‘macrobes’, which explains why environmental filtering often is the dominant process in microbial community assembly (Van der Gucht *et al.*, 2007, Martiny *et al.*, 2011, Lindstrom & Langenheder, 2012, Zhang *et al.*, 2019). Our results on dispersal limitation of the five community types reflect the propagule size of the organisms: seeds are larger than fungal spores that are larger than bacterial cells. This suggests that airborne dispersal is mostly shaping the distribution of bacteria, fungi, and plants in our study system. Next to the dispersal abilities of the organisms, the exposure of the habitats to the environment can determine whether dispersal is shaping community assembly. Leaves are more exposed to long-distance dispersed microbes than soil that is less exposed to wind and rain, which may explain why soil microbial communities appeared to be more dispersal limited than leaf communities.

Soil microbial communities and plants completely depend on the water and nutrient availability as well as on chemical and physical properties of the soil they are living on, which is reflected by the strong observed environmental filtering in these communities. Many of these soil properties are modified by the plant species using the soil as substrate (Bulgarelli *et al.*, 2013) explaining why soil microbial communities strongly responded to the plant communities on the plots. Leaf-associated microbes live in their own environment characterized by low availability of nutrients and water, plant metabolites, and strong radiation (Vorholt, 2012), which is buffered from the environmental conditions experienced by soil microbes and plants. Accordingly, not the environmental conditions on the plot but plant-species identity, which is a proxy for differences in leaf properties and thus the niches provided by leaves, strongly affected the composition of the leaf microbiome.

All community types appeared to be more segregated than expected by chance, which may indicate that antagonistic interactions such as competition or inhibition are the dominant factor in the species interactions in the communities observed here. Recently Blanchet *et al.* (2020) discussed that co-occurrence patterns are poor proxies for species interactions. For instance, shared or exclusive niches may lead to aggregation or segregation, respectively, independently of direct interferences between organisms. Particularly in microbial communities where interactions are often restricted to other microbes in direct proximity (Cordero & Datta, 2016, Dal Co *et al.*, 2020), co-occurrence patterns may be not indicative for interactions also because one sample integrates over a number of niches. Even though these results may not be conclusive for the type and strength of interactions, the fact that all community types were more segregated than expected by chance suggests that at least strong facilitation is not common in these communities. Finally, soil seems not to be a major source for microbes associated with leaves although some microbial strains were found in both soil and leaf samples. We collected bulk soil and not specifically the rhizosphere of individual species, which may have caused the low overlap of soil and leaf microbes in our samples.

The approach to dissect community assembly into separated processes or ‘filters’ that act hierarchically on the regional species pool and shape local species assemblage has been criticized (Gilbert & Bennett, 2010, Stegen & Hurlbert, 2011, Kraft *et al.*, 2015, Stegen *et al.*, 2015, Cadotte & Tucker, 2017, Blanchet *et al.*, 2020). As detailed by these authors, the outcomes of our predictions deduced from assembly hypotheses may be the result from the process under consideration, or the result from another process that is overlaying the other one. Shared niches among species will lead to a strong signal in the environmental and the interaction filter. *Vice versa,* strong mutualistic interactions leading to high co-occurrence may be misinterpreted as result of environmental filtering. Furthermore, in our study site the effect of the environmental and the dispersal filter do not act independently as abiotic conditions and geographic distance covary along a successional gradient. Finally, composition data alone may not carry sufficient information to infer assembly processes. Thus, the observed pattern for soil microbes and plants cannot be attributed specifically to one of these filters, but most likely they jointly contribute to the findings reported here. Therefore, our analysis may not be suitable to identify individual processes that dominate the assembly of one of the communities observed. However, our comparative approach considering different organisms that live on different substrates but within the same landscape and recorded in close proximity in the same plots is well suited to highlight the characteristics of each of the community types and how this affects their assembly. In summary, the differences in community assembly processes can be attributed to the size of the organisms’ propagules and thus their dispersal abilities, the exposure of the habitats to the environment (i.e., the accessibility to airborne propagules), and the characteristics of the habitat itself (i.e., soil *versus* leaves). Additionally, the spatial scale and thus the heterogeneity of niches within a sample affect most of the processes discussed in this study: Microbial propagules experience less dispersal limitations than plant propagules on the landscape scale. However, a few millimetres in the soil or on leaves may represent a strong barrier for resident microbes that rely on specific niches. Finally, as discussed above, a one-square-meter-plot hosts a community of potentially interacting plants within a shared niche; a soil of leaf sample contains multiple niches with distinct microbial communities. Our study identified organismal traits and abiotic factors that may affect community assembly and may thus stimulate further work on assembly processes conditional to the characteristics of the organism, the habitat, and the spatial scale under consideration.

## Acknowledgements

We thank the Hohe Tauern National Park Salzburg Administration and the Rudolfshütte for organizational and logistic support, the governing authority Land Salzburg for the permit to conduct our research (permit # 20507-96/45/7-2019). Hamed Azarbad and Lisa-Maria Ohler provided valuable comments to an earlier version of the manuscript. The study was supported by the START-program of the Austrian Science Fund (FWF) granted to RRJ (Y1102).

## Supplementary Information

### SI1 Summary and details of statistical results

### Supporting Information 1

#### H1: dispersal filter

Mantel statistic based on Pearson’s product-moment correlation

|  | Sample | r | p |
| --- | --- | --- | --- |
| Bacteria | plants (all species) | 0.059 | 0.002 |
| Fungi | plants (all species) | 0.146 | 0.001 |
| Plant | NA | 0.452 | 0.001 |
| Bacteria | soil | 0.376 | 0.001 |
| Fungi | soil | 0.396 | 0.001 |
| Bacteria | *Campanula scheuchzeri* | 0.159 | 0.001 |
| Fungi | *Campanula scheuchzeri* | 0.205 | 0.001 |
| Bacteria | *Oxyria digyna* | -0.007 | 0.504 |
| Fungi | *Oxyria digyna* | 0.152 | 0.047 |
| Bacteria | *Trifolium badium* | 0.014 | 0.396 |
| Fungi | *Trifolium badium* | -0.011 | 0.521 |

#### H2: environmental filter

Mantel statistic based on Pearson’s product-moment correlation

H2: environmental filter Mantel statistic based on Pearson's product-moment correlation
| Abiotic factors |  |  |  |
| --- | --- | --- | --- |
|  | Sample | r | p |
| Bacteria | plants (all species) | -0.01626 | 0.744 |
| Fungi | plants (all species) | 0.1257 | 0.002 |
| Plant | NA | 0.4584 | 0.001 |
| Bacteria | soil | 0.3319 | 0.001 |
| Fungi | soil | 0.3638 | 0.001 |
| Bacteria | *Campanula scheuchzeri* | -0.09257 | 0.939 |
| Fungi | *Campanula scheuchzeri* | 0.06553 | 0.169 |
| Bacteria | *Oxyria digyna* | 0.1274 | 0.178 |
| Fungi | *Oxyria digyna* | 0.09107 | 0.213 |
| Bacteria | *Trifolium badium* | -0.1109 | 0.877 |
| Fungi | *Trifolium badium* | 0.1806 | 0.109 |

| Plant community as environment |  |  |  |
| --- | --- | --- | --- |
|  | Sample | r | p |
| Bacteria | plants (all species) | 0.03252 | 0.048 |
| Fungi | plants (all species) | 0.1967 | 0.001 |
| Bacteria | soil | 0.61 | 0.001 |
| Fungi | soil | 0.6206 | 0.001 |
| Bacteria | *Campanula scheuchzeri* | 0.0691 | 0.074 |
| Fungi | *Campanula scheuchzeri* | 0.2718 | 0.001 |
| Bacteria | *Oxyria digyna* | 0.1384 | 0.068 |
| Fungi | *Oxyria digyna* | 0.148 | 0.022 |
| Bacteria | *Trifolium badium* | -0.08512 | 0.86 |
| Fungi | *Trifolium badium* | 0.3236 | 0.015 |

#### H3: environmental filter

Distance-based redundancy analyses using Bray-Curtis distances followed by permutation test under reduced model

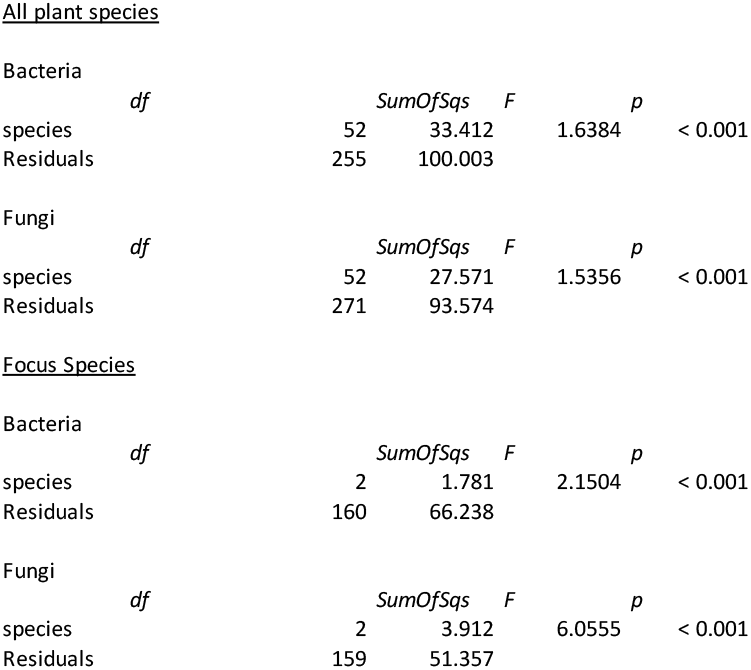

#### H4: interaction filter

cooc_null_model function implemented in the R package EcoSimR

bacteria

*Campanula scheuchzeri*

Time Stamp: Mon Oct 26 16:38:55 2020

Reproducible:

Number of Replications:

Elapsed Time: 14 mins

Metric: c_score

Algorithm: sim9

Observed Index: 2.4138

Mean Of Simulated Index: 2.41

Variance Of Simulated Index: 7.0751e-07

Lower 95% (1-tail): 2.4084

Upper 95% (1-tail): 2.4114

Lower 95% (2-tail): 2.4084

Upper 95% (2-tail): 2.4116

Lower-tail P > 0.999

Upper-tail P < 0.001

Observed metric > 1000 simulated metrics

Observed metric < 0 simulated metrics

Observed metric = 0 simulated metrics

Standardized Effect Size (SES): 4.6055

bacteria

*Oxyria digyna*

Time Stamp: Mon Oct 26 16:47:30 2020

Reproducible:

Number of Replications:

Elapsed Time: 42 secs

Metric: c_score

Algorithm: sim9

Observed Index: 1.4072

Mean Of Simulated Index: 1.398

Variance Of Simulated Index: 2.6967e-06

Lower 95% (1-tail): 1.3957

Upper 95% (1-tail): 1.4007

Lower 95% (2-tail): 1.3953

Upper 95% (2-tail): 1.4009

Lower-tail P > 0.999

Upper-tail P < 0.001

Observed metric > 1000 simulated metrics

Observed metric < 0 simulated metrics

Observed metric = 0 simulated metrics

Standardized Effect Size (SES): 5.5786

bacteria

*Trifolium badium*

> summary(myModel)

Time Stamp: Mon Oct 26 17:04:55 2020

Reproducible:

Number of Replications:

Elapsed Time: 15 mins

Metric: c_score

Algorithm: sim9

Observed Index: 1.7374

Mean Of Simulated Index: 1.7358

Variance Of Simulated Index: 1.4354e-07

Lower 95% (1-tail): 1.7352

Upper 95% (1-tail): 1.7364

Lower 95% (2-tail): 1.7351

Upper 95% (2-tail): 1.7364

Lower-tail P > 0.999

Upper-tail P < 0.001

Observed metric > 1000 simulated metrics

Observed metric < 0 simulated metrics

Observed metric = 0 simulated metrics

Standardized Effect Size (SES): 4.2524

bacteria

Soil

Time Stamp: Sun Nov 1 10:39:07 2020

Reproducible:

Number of Replications:

Elapsed Time: 49 mins

Metric: c_score

Algorithm: sim9

Observed Index: 8.1985

Mean Of Simulated Index: 8.1884

Variance Of Simulated Index: 6.2715e-06

Lower 95% (1-tail): 8.1847

Upper 95% (1-tail): 8.1932

Lower 95% (2-tail): 8.1845

Upper 95% (2-tail): 8.1934

Lower-tail P > 0.999

Upper-tail P < 0.001

Observed metric > 1000 simulated metrics

Observed metric < 0 simulated metrics

Observed metric = 0 simulated metrics

Standardized Effect Size (SES): 4.0201

fungi

*Campanula scheuchzeri*

summary(myModel)

Time Stamp: Thu Feb 4 17:23:12 2021

Reproducible:

Number of Replications:

Elapsed Time: 4.2 mins

Metric: c_score

Algorithm: sim9

Observed Index: 7.3166

Mean Of Simulated Index: 7.2955

Variance Of Simulated Index: 0.00012776

Lower 95% (1-tail): 7.2795

Upper 95% (1-tail): 7.3105

Lower 95% (2-tail): 7.2785

Upper 95% (2-tail): 7.3106

Lower-tail P > 0.999

Upper-tail P < 0.001

Observed metric > 1000 simulated metrics

Observed metric < 0 simulated metrics

Observed metric = 0 simulated metrics

Standardized Effect Size (SES): 1.8662

fungi

*Oxyria digyna*

Time Stamp: Fri Feb 5 08:34:09 2021

Reproducible:

Number of Replications:

Elapsed Time: 36 secs

Metric: c_score

Algorithm: sim9

Observed Index: 1.6337

Mean Of Simulated Index: 1.6016

Variance Of Simulated Index: 6.983e-05

Lower 95% (1-tail): 1.5922

Upper 95% (1-tail): 1.6153

Lower 95% (2-tail): 1.5918

Upper 95% (2-tail): 1.6157

Lower-tail P > 0.999

Upper-tail P < 0.001

Observed metric > 1000 simulated metrics

Observed metric < 0 simulated metrics

Observed metric = 0 simulated metrics

Standardized Effect Size (SES): 3.8358

fungi

*Trifolium badium*

Time Stamp: Fri Feb 5 08:34:58 2021

Reproducible:

Number of Replications:

Elapsed Time: 3.5 secs

Metric: c_score

Algorithm: sim9

Observed Index: 1.6781

Mean Of Simulated Index: 1.6312

Variance Of Simulated Index: 0.00019536

Lower 95% (1-tail): 1.6065

Upper 95% (1-tail): 1.6502

Lower 95% (2-tail): 1.6063

Upper 95% (2-tail): 1.6512

Lower-tail P > 0.999

Upper-tail P < 0.001

Observed metric > 1000 simulated metrics

Observed metric < 0 simulated metrics

Observed metric = 0 simulated metrics

Standardized Effect Size (SES): 3.3565

fungi

Soil

Time Stamp: Mon Nov 2 16:58:41 2020

Reproducible:

Number of Replications:

Elapsed Time: 58 mins

Metric: c_score

Algorithm: sim9

Observed Index: 3.7107

Mean Of Simulated Index: 3.7092

Variance Of Simulated Index: 1.1674e-07

Lower 95% (1-tail): 3.7088

Upper 95% (1-tail): 3.7099

Lower 95% (2-tail): 3.7088

Upper 95% (2-tail): 3.7099

Lower-tail P > 0.999

Upper-tail P < 0.001

Observed metric > 1000 simulated metrics

Observed metric < 0 simulated metrics

Observed metric = 0 simulated metrics

Standardized Effect Size (SES): 4.379

plants

Time Stamp: Thu Oct 29 14:24:04 2020

Reproducible:

Number of Replications:

Elapsed Time: 2.1 secs

Metric: c_score

Algorithm: sim9

Observed Index: 148.88

Mean Of Simulated Index: 144.45

Variance Of Simulated Index: 0.057293

Lower 95% (1-tail): 144.03

Upper 95% (1-tail): 144.87

Lower 95% (2-tail): 143.99

Upper 95% (2-tail): 144.91

Lower-tail P > 0.999

Upper-tail P < 0.001

Observed metric > 1000 simulated metrics

Observed metric < 0 simulated metrics

Observed metric = 0 simulated metrics

Standardized Effect Size (SES): 18.484

#### H5: subset hypothesis

|  |  | proportion |
| --- | --- | --- |
| H5 | Bacteria | 0.03587705 |
| H5 | Fungi | 0.05591168 |

